# Skeletal trait measurements for thousands of bird species

**DOI:** 10.1101/2024.12.19.629481

**Authors:** Brian C. Weeks, Zhizhuo Zhou, Charlotte M. Probst, Jacob S. Berv, Bruce O’Brien, Brett W. Benz, Heather R. Skeen, Mark Ziebell, Louise Bodt, David F. Fouhey

**Author notes:** Corresponding author: Brian Weeks.

## Abstract

Large comparative datasets of avian functional traits have been used to address a wide range of questions in ecology and evolution. To date, this work has been constrained by the limited availability of skeletal trait datasets that include extensive inter- and intra-specific sampling. We use computer vision to identify and measure bones from photographs of museum skeletal specimens to assemble an extensive dataset of functionally important skeletal elements in birds. The dataset spans 2,057 species of birds (Aves: Passeriformes) and includes measurements of 12 skeletal elements from 14,419 individuals. In addition to the trait values directly measured from photographs, we leverage the multi-dimensional nature of our dataset and known phylogenetic relationships of the species to impute missing data under an evolutionary model. To facilitate use of the dataset, the taxonomy has been reconciled with an existing comprehensive avian phylogeny and an additional dataset of external functional traits for all birds.

## Background & Summary

Understanding large-scale patterns of diversity, and the ecological and evolutionary origins and consequences of these patterns, is of growing interest. These efforts have historically been constrained by the limited availability of comparative quantitative trait datasets at large spatial and taxonomic scales^1^. As large datasets have become available, they have stimulated significant advances in macroecology and macroevolution.

While large-scale trait datasets are increasingly available across a range of taxa (e.g., vascular plants^2^, lizards^3^, and freshwater fish^4^), birds are a model system in macroecology and macroevolution, with their well-known distributions^5^, extinction risks^5^, phylogenetic relationships^6,7^, ecological niches^8,9^, life history strategies^10^, nesting biologies^11^, and external morphologies^12^. These diverse datasets have been integrated to answer a wide range of questions spanning ecology and evolution (e.g.^13–17^) and are increasingly being used to understand human impacts on natural systems (e.g.^18–22^). Although much has been learned from existing large-scale datasets, in animals, the availability of trait data spanning multiple anatomical systems would open new avenues of research and could allow for more mechanistic understanding of morphological patterns. Bird skeletons, which are well-represented in natural history collections, present an underutilized opportunity to develop such a dataset.

The accumulation of comparative skeletal trait data for many traits and across many species and individuals has lagged far behind the generation of data from the measurement of external traits^23^. In birds, aspects of the skeleton provide key insights into bird locomotion^24^, the physics of flight^25,26^, directional evolution^27^, phylogenetic relationships^28^, and responses to environmental change^29^, and are often used to better understand the morphologies of fossil birds^30^. Further, the utility of skeletal traits expands significantly when they can be studied in conjunction with other types of phenotypic data. For example, although the tendency for appendages to be longer in warmer climates (i.e., Allen’s Rule^31^) is a classic pattern in macroecology and has been the focus of intensive research for over a century, the integration of skeletal and plumage trait data revealed a novel morphological trend that generated new insights into the mechanistic basis underlying Allen’s Rule^32^. As such, the availability of comparative skeletal data may provide novel insight into macro-scale patterns in avian morphology and improved mechanistic understanding of the drivers of bird morphological variation across space and time.

This dataset encompasses a large portion of the diversity within Passeriformes, the most diverse order of Neornithes (modern, living birds). The 2,057 species in the dataset comprise 34% of passerine species and represent 89% of passerine families^6^. The sampling is also spatially expansive and includes specimens from all continents except Antarctica (Fig 1). Multiple individuals were measured per species when possible, resulting in a dataset that includes 14,419 individuals. We targeted twelve skeletal elements for each specimen. Combined with our estimates for missing values our dataset includes 173,028 unique values. The data are presented in three formats: 1) a specimen-level dataset that only includes trait values that were directly measured, 2) a specimen-level dataset with no missing data that includes both the directly measured trait values and imputed trait values, and 3) a complete species-level dataset derived by applying a multivariate evolutionary model. The taxonomy in the datasets has been unified to the Birdlife Version 3 taxonomy to facilitate integration with existing largescale datasets and to simplify conversion to other widely used taxonomies using recently published taxonomic crosswalks^12^. As such, it should be straightforward to integrate our data with data on the phylogenetic history of birds^6^, bird range maps^5^, IUCN threat statuses^5^, and existing comprehensive external trait data^12^. Importantly, the methods used to generate these data are open source^23^ and easily applied, enabling future expansion of the dataset.

**Figure 1.**
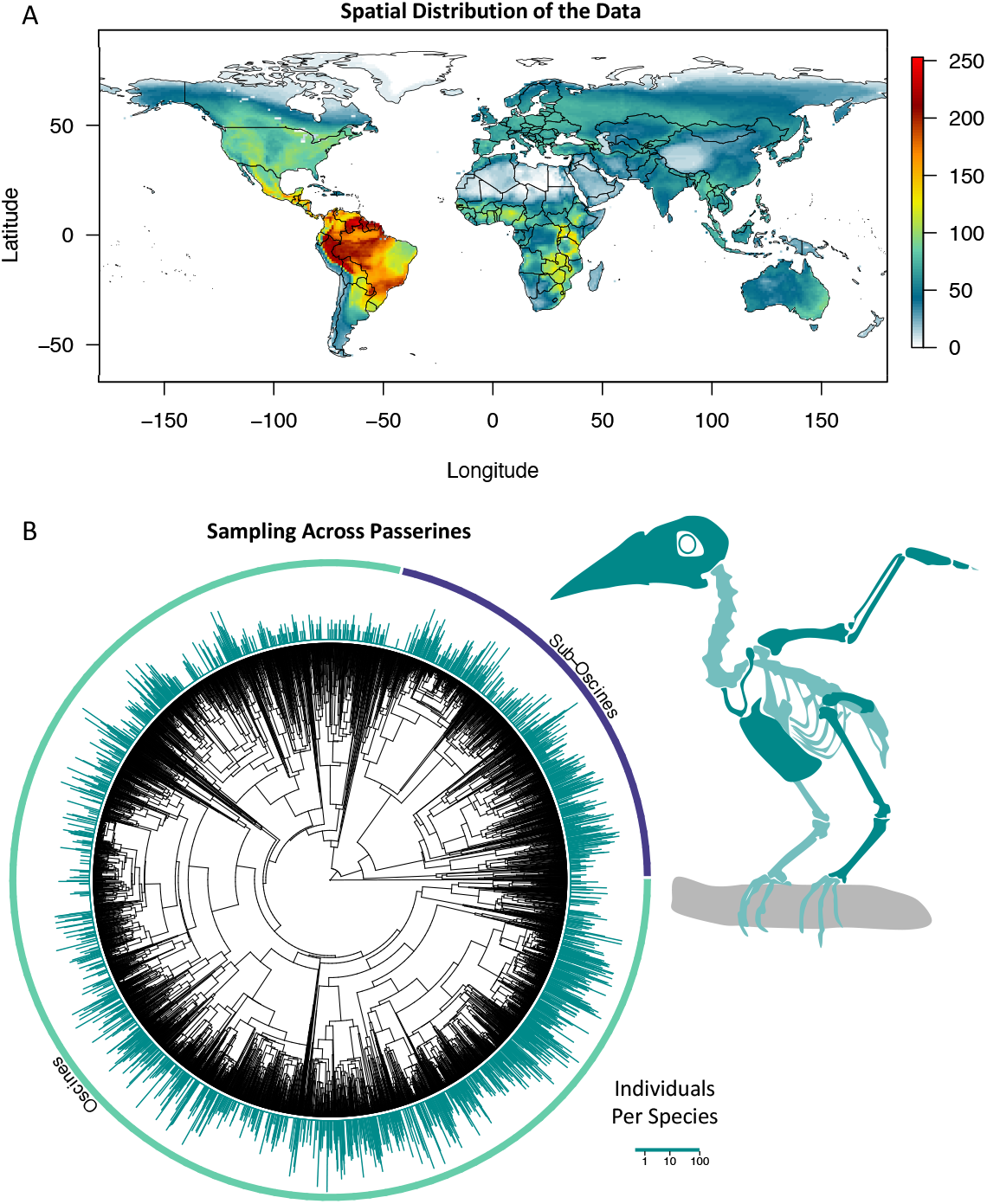
Dataset coverage. (**A**) The dataset includes species spanning all continents except Antarctica. The ranges of all species included in the dataset are plotted, with colour indicating the number of species included in our dataset at each point in space. (**B**) The species in the dataset span Passeriformes, the most diverse taxonomic order within modern birds. Each bar associated with a tip on the phylogeny represents a species that is included in the dataset, with the height of the bar indicating the number of individuals of that species that were measured and included in the dataset. The bird skeleton highlights the bones that were measured in darker green.

## Methods

### Sampling

The large majority of our data come from skeleton specimens held in the University of Michigan Museum of Zoology (UMMZ), one of the largest and most diverse bird skeletal collections in the world, where we effectively photographed and measured the entirety of the UMMZ’s passerine skeletal collection (*N* = 12,421, number of species = 1,881). We supplemented this dataset with specimens at the Field Museum of Natural History (FMNH; *N* = 1,998, number of species = 438), targeting families that are well represented in both the UMMZ and FMNH collections, with an emphasis on species found in the Neotropics. Ultimately, we photographed 14,419 specimens spanning 2,057 species, from 86 families (Fig. 1). Every trait measurement has an associated specimen catalogue number that can be used to link each measurement to the specimen, housed in UMMZ or FMNH.

### Photographing

Trait measurements were generated using Skelevision^23^, a deep neural network-based approach for identifying and measuring skeletal elements in photographs of bird skeleton specimens. In this approach, museum skeleton specimens are first removed from their containers and spread randomly on a standard background, except for the keel and the skull, which are consistently oriented to display their profile. They are then photographed from a fixed distance before being returned to their boxes. Each specimen is photographed individually, independently, and in its entirety.

The photographs were taken with the same imaging equipment that was used in the Skelevision methods paper^23^. All images were collected from ∼400mm above the specimen, using a SONY IX183 sensor on a FLIR Blackfly S camera. This generated photographs with a pixel size of 0.07mm.

### Trait Measurement

We applied the Skelevision method^23^ for segmenting, identifying, and measuring target bones to each photograph. This method integrates a U-Net^33^ and Mask R-CNN^34^ trained on images annotated by hand^35,36^ to identify pixels in the images that are bone, determine which element the pixels belong to, and then segment the elements. The pipeline then takes segmented masks from the images for all elements and measures their longest linear dimension (i.e., the longest linear length of each element)^23^. We use this method to measure 12 traits: the lengths of the tibiotarsus, humerus, tarsometatarsus, ulna, radius, keel, carpometacarpus, 2^nd^ digit 1^st^ phalanx, furcula, and femur; the maximum outer diameter of the sclerotic ring, and the length from the back of the skull to the tip of the bill (treating the rhamphotheca as part of the bill when it remains present on the specimen).

Skelevision estimates the probability that Skelevision’s classification of each element is correct, given the classification options (‘bprob’). Because elements that are classified with a lower certainty (i.e., a low bprob) are at a higher risk of false positives^23^, we filtered out all trait estimates with a bprob < 0.95; this has been found to result in a relatively low rate of false negatives without increasing the risk of false positives^23^. For specimens with multiple high-confidence estimates of a trait (e.g., if two femurs were confidently identified and measured from a specimen), we combined these measures by taking the mean. In this way, a single high-quality estimate of each trait was made for each specimen whenever at least one example of an element was confidently identified (*Skelevision-Only Dataset*). For those traits that did not have at least one high confidence trait estimate (i.e., if there was not at least one trait measure with a bprob ≥ 0.95), the trait was marked as missing data (given a value of ‘NA’ in the *Skelevision-Only Dataset*).

### Phylogenetic Data Imputation and Validation

To generate a 100% complete dataset, we imputed values for all missing data in the *Skelevision-Only Dataset* using Rphylopars, a maximum likelihood approach for fitting multivariate phylogenetic models and estimating missing values in comparative data^37^. An advantage of this approach is that it can model variation at the level of individual specimens along with variation among species, providing estimates of missing values at the individual level. This approach allowed us to leverage the dataset’s large size and dimensionality (12 dimensions x 14,419 individuals), along with the expected hierarchical structure due to phylogeny, to estimate missing values. To approximate phylogenetic relationships among the included species, we downloaded 1,000 trees from the posterior distribution of a phylogeny for all birds^6,38^ and constructed a consensus tree following Rubolini *et al*.^39^. Using this consensus tree, we used Rphylopars to estimate variance-covariance structures (both within and between species) according to the expectations of a multivariate Brownian Motion model (‘mvBM’) process. All data were log-transformed before model fitting with Rphylopars, but otherwise, we left all user options set at their defaults (see *Code Availability* for details on accessing the ‘Trait_Imputation.R’ script).

To evaluate the accuracy of trait imputation, we validated our approach by iteratively withholding 10-90% of the non-missing data for each trait to use as test data. We then estimated the mvBM model, excluding the test data in the model fitting, and imputed the withheld test data. For each level of missing data, we repeated the analyses ten times, selecting a different random sample of the dataset to use as test data for each replicate. We then compared the known true values of the test data to their estimated values by calculating the root-mean-squared error (RMSE) and percentage bias (p-bias; Fig. 2).

**Figure 2.**
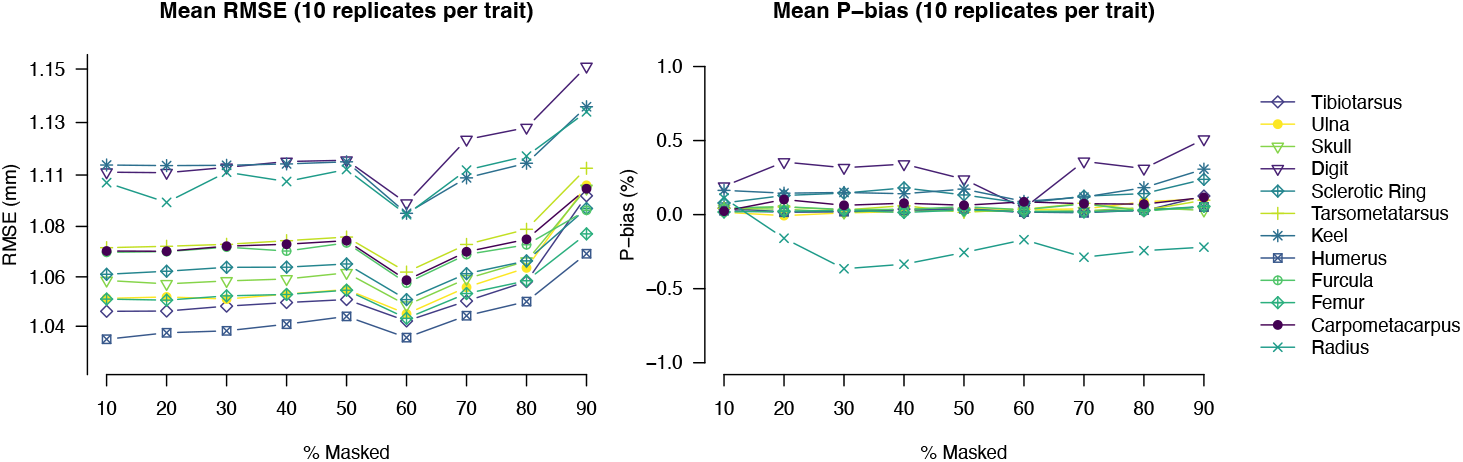
Variation in trait imputation accuracy across different levels of missing data. We masked an increasing percentage of the data generated by Skelevision (10-90% in intervals of 10%) and then imputed the missing values. We estimate the RMSE and P-Bias across ten randomized sets of data for each percentage quantile and present the mean per trait. RMSE and P-Bias are generally low and stable across the range of missing data quantiles.

We present the data from these analyses as complete datasets both at the specimen level (*Complete Trait Dataset*) and at the species level using species averages estimated by Rphylopars, which includes both the averages and the estimated standard error associated with each species mean (*Species-level Data Estimates*).

## Data Records

For all datasets, we include species binomials following the BirdLifeV3 taxonomy. For the specimen-level data, we also include museum specimen catalogue numbers. The data are presented in three datasets:

### Skelevision-Only Dataset

This dataset includes all the Skelevision data measured with high confidence (bprob ≥ 0.95). It does not include any imputed trait measurements. The data are available for download as a comma-separated values file, “Skelevision_Only_Dataset_v1.csv”. DOI to be provided following peer review.

### Complete Trait Dataset

This dataset includes all the Skelevision data measured with a high degree of confidence and imputed trait values for all missing data. The data are available for download as a comma-separated values file, “Complete_Trait_Dataset_v1.csv”. DOI to be provided following peer review.

### Species-level Data Estimates

This dataset includes species means from the model fit using Rphylopars. We also provide estimates of the species-level standard error, variance, and 95% confidence intervals around trait means for downstream analyses. The data are available for download as a comma-separated values file, “Skelevision_species_complete_v1”. DOI to be provided following peer review.

## Technical Validation

### Skelevision Accuracy

The accuracy of the processed data (*Skelevision-Only Dataset*) generated from the UMMZ image capture setup and specimens has been quantified previously. Weeks et al (2023) compared 100 handmade measurements of each trait (except the furcula) to Skelevision estimates of the same traits on the same specimen and found a mean RMSE of 0.89mm, with some variation in error across bone types (Table 1).

**Table 1.** Estimated error of Skelevision-generated data. Data for 100 specimens photographed at UMMZ were compared to measurements made by a single measurer, who measured the same bone by hand using digital callipers. The root mean square error (RMSE) and percent bias (P-bias) of the Skelevision estimates compared to the handmade measurements, taken from^23^, are presented. For 30 FMNH specimens, a single measurer used ImageJ software^40^ to measure each of the traits in the photographs; as with the UMMZ specimens, we present the RMSE and P-bias for the Skelevision estimates compared to the handmade estimates from the same specimen.

| Skeletal Element | UMMZ-Human<br>RMSE<sup>†</sup> | UMMZ-Human<br>P-bias<sup>†</sup> | FMNH-Human<br>RMSE | FMNH-Human<br>P-bias |
| --- | --- | --- | --- | --- |
| Skull | 1.20 mm | -1.8% | 0.68 mm | -0.7% |
| Sclerotic Ring | 0.34 mm | 2.4% | 0.25 mm | 1.0% |
| Humerus | 0.47 mm | -0.2% | 0.53 mm | 2.3% |
| Radius | 0.51 mm | -0.5% | 1.39 mm | -0.7% |
| Ulna | 0.96 mm | -1.6% | 0.86 mm | -0.6% |
| Carpometacarpus | 1.11 mm | 0.5% | 1.64 mm | 1.3% |
| 2<sup>nd</sup> Digit 1<sup>st</sup><br>Phalanx | 0.49 mm | 3.3% | 3.14 mm | 20.4% |
| Furcula | NA | NA | 3.31 mm | 4.0% |
| Keel | 2.23 mm | -6.1% | 4.63 mm | 8.5% |
| Femur | 0.46 mm | 0.0% | 0.39 mm | 0.6% |
| Tibiotarsus | 1.17 mm | -0.7% | 0.77 mm | 2.1% |
| Tarsometatarsus | 0.84 mm | 0.6% | 3.74 mm | 9.6% |

<sup>†</sup>Estimates for the UMMZ-human RMSE and P-bias are taken from <sup>23</sup>

Because there is a risk that Skelevision will perform differently across the different image-capturing contexts (e.g., locations with variation in lighting) and some variation in specimen preparation between UMMZ and FMNH (e.g., differences in the degree to which bones remain articulated), and to collect validation data for the furcula, we conducted a similar validation test with a subset of the FMNH data. For a random sample of 30 specimens from FMNH, a single person measured each trait of interest from the photographs of the specimens using ImageJ software^40^. We then compared these handmade measurements of the trait values to the Skelevision measurements for the same trait on the same skeleton. As with the UMMZ samples, we find Skelevision is accurate, with a mean RMSE of 1.78 mm across all traits. This is higher than the RMSE of the UMMZ specimens but remains comparable to inter-human measurement error. The errors are not uniform among the element types, and while many have similar or lower errors compared to the UMMZ data, a few have elevated error levels, albeit on a similar scale to the expected range of human measurement error (Table 1)^9^.

### Trait Imputation Accuracy

The RMSE of the imputed data was uniformly low across replicates and increasing levels of additional missing data (Table 2; Figure 2). This suggests potential uncertainty in phylogenetic relationships has a negligible impact on the trait imputation accuracy. The maximum mean RMSE (∼ 1.15mm) was observed for estimates of the length of the second digit; notably this was only when we simulated maximal amounts of additional missing data (90%; or ∼85% missing data overall). This maximum mean error is like that observed from human measurement^9^. We also observe very low P-bias (< 1%) in estimated values throughout the range of evaluated levels of missing data, lending further credence to the validity of our approach. Maximum P-bias values ranged from ∼ -0.4% (radius) to 0.5% (second digit) and thus were essentially negligible, particularly at the levels of missing data within the dataset (Table 2); most traits had a mean P-bias centered near zero.

**Table 2.** Estimated error of imputed data. We round up the percent missing data to the nearest ten and include the predicted RMSE and P-bias for that trait at that level of missing data (Fig. 2).

| Skeletal Element | Specimens<br>with Direct<br>Measurements | Percent<br>Missing Data | Predicted RMSE | Predicted P-bias |
| --- | --- | --- | --- | --- |
| Skull | 13,197 | 8% | 1.06 mm | 0.04% |
| Sclerotic Ring | 8,703 | 40% | 1.07 mm | 0.18% |
| Humerus | 12,306 | 15% | 1.04 mm | 0.02% |
| Radius | 7,274 | 50% | 1.11 mm | -0.26% |
| Ulna | 8,574 | 41% | 1.06 mm | 0.04% |
| Carpometacarpus | 7,681 | 47% | 1.08 mm | 0.06% |
| 2<sup>nd</sup> Digit 1<sup>st</sup><br>Phalanx | 4,635 | 68% | 1.12 mm | 0.36% |
| Furcula | 5,473 | 62% | 1.07 mm | 0.07% |
| Keel | 10,842 | 25% | 1.11 mm | 0.15% |
| Femur | 12,584 | 13% | 1.05 mm | 0.02% |
| Tibiotarsus | 12,579 | 13% | 1.05 mm | 0.02% |
| Tarsometatarsus | 8,921 | 38% | 1.08 mm | 0.06% |

In general, when anatomical traits have strong phylogenetic signals and multivariate correlations, we expect to be able to estimate missing values with high accuracy and precision under mvBM. Our approach highlights the power of multivariate phylogenetic models to generate complete datasets at the level of individual specimens and should provide a useful framework for future research.

## Code Availability

The code to generate Skelevision trait estimates is publicly available (DOI: 10.5281/zenodo.6402893). We have also provided all the code used to impute the trait data, ‘Trait_Imputation.R’, with the data on Dryad (DOI to be provided following peer review). The validation data and the script used to quantify RMSE are also included with the data on Dryad (DOI to be provided following peer review). All trait imputation was done in R^41^. Images of all UMMZ specimens are available through the Deep Blue Data Repository (DOI to be provided following peer review).

## Acknowledgements

We thank Benjamin Winger, Bird Division, University of Michigan Museum of Zoology for granting us access to the UMMZ specimens. We thank John Bates, Shannon Hackett, and Benjamin Marks, Division of Birds, Field Museum of Natural History, for granting us access to the FMNH specimens. This work was supported by the David and Lucile Packard Foundation. JSB was supported by an Eric and Wendy Schmidt AI in Science Postdoctoral Fellowship, Schmidt Futures.

## Author contributions

BCW: Conceptualization, Methodology, Validation, Formal analysis, Investigation, Resources, Data Curation, Project Administration, Writing – Original Draft

ZZ: Methodology, Formal analysis, Investigation, Writing – Review and Editing

CP: Methodology, Validation, Investigation, Writing – Review and Editing

JB: Methodology, Validation, Formal analysis, Writing – Original

Draft BO: Investigation, Writing – Review and Editing

BB: Project Administration, Writing – Review and Editing

HS: Investigation, Writing – Review and Editing

MZ: Investigation, Writing – Review and Editing

LB: Investigation, Writing – Review and Editing

DFF: Conceptualization, Methodology, Formal Analysis, Resources, Project Administration, Writing – Review and Editing

## Competing interests

The authors declare no competing interests.

## Notes

### Competing Interest Statement

The authors have declared no competing interest.

